# Brassinosteroid controls leaf air space patterning non-cell autonomously by promoting epidermal growth

**DOI:** 10.1101/2025.07.14.664710

**Authors:** James M. Fitzsimons, Ana B. Rock, Richard L. De Falbe, Samantha Fox, Chris D. Whitewoods

## Abstract

Plant leaves contain a complex network of intercellular air spaces, which enhance gas exchange and allow efficient photosynthesis. However, despite their importance, how leaf air spaces are patterned is poorly understood. It has been proposed for almost a century that air spaces form by faster-growing epidermal tissue pulling slower-growing mesophyll cells apart, but this has never been tested. Here we characterise air space morphogenesis throughout the entirety of leaf development in the first leaf of *A. thaliana* and show that the plant hormone brassinosteroid (BR) is required for air space expansion in the palisade, but not the spongy, mesophyll. We also show that epidermal brassinosteroid perception is sufficient to promote air space expansion in the palisade non cell-autonomously and propose that this non cell-autonomous effect is due to altered epidermal growth. To test if epidermal growth affects air space patterning we reduce growth specifically in the epidermis using inducible expression of the growth repressor BIG BROTHER and show that an epidermal growth restriction reduces air space expansion in the palisade mesophyll. Overall, we propose that brassinosteroid signalling promotes growth in the epidermis to pattern air spaces in the palisade mesophyll.

**Summary statement:** Up to 70% of leaf volume is intercellular air space. This work shows that the plant hormone brassinosteroid acts in the epidermis to promote air space morphogenesis non-cell autonomously.

## Introduction

The size and complexity of organisms are limited by their ability to obtain nutrients from their environment by diffusion (Haldane, 1927). To overcome this limitation, multicellular animals and plants have evolved internal structures with large surface areas to increase nutrient absorption. The convoluted air spaces found within the lungs of animals and the leaves of plants are key examples of this. Air spaces bring internal cells into contact with the atmosphere and mediate the efficient exchange of carbon dioxide and oxygen gases necessary for respiration and photosynthesis. In animals, the developmental and molecular mechanisms of air space formation within lungs are well studied, with several genes interacting to control elongation and branching of an air-filled tissue tube (Metzger *et al*., 2008; Nikolić, Sun and Rawlins, 2018). However, in plants air space development is poorly understood. This is all the more striking as air spaces form up to 70% of leaf volume (Earles *et al*., 2018) and affect leaf photosynthetic efficiency (Lehmeier *et al*., 2017; Mathers *et al*., 2018).

Leaf air spaces form by cell separation at multicellular junctions (Jeffree, Dale and Fry, 1986; Heynh *et al*., 1991; Kalve *et al*., 2014), rather than by cell death as in roots of many species (Evans, 2004). These smaller air spaces expand and join to form an interconnected network throughout the leaf, patterned precisely in three dimensions. The upper half of many leaves contains densely packed palisade mesophyll cells with only small air spaces (fig. 1 A and B) whereas the lower half contains lobed spongy mesophyll cells with larger air spaces (fig. 1 A and C). The largest spaces are positioned adjacent to stomata. This arrangement is thought to maximise light harvesting in the top half (Terashima and Inoue, 1984) and gas exchange and light scattering in the lower half to increase overall photosynthetic efficiency (DeLucia *et al*., 1996; Smith *et al*., 1997).

**Figure 1.**
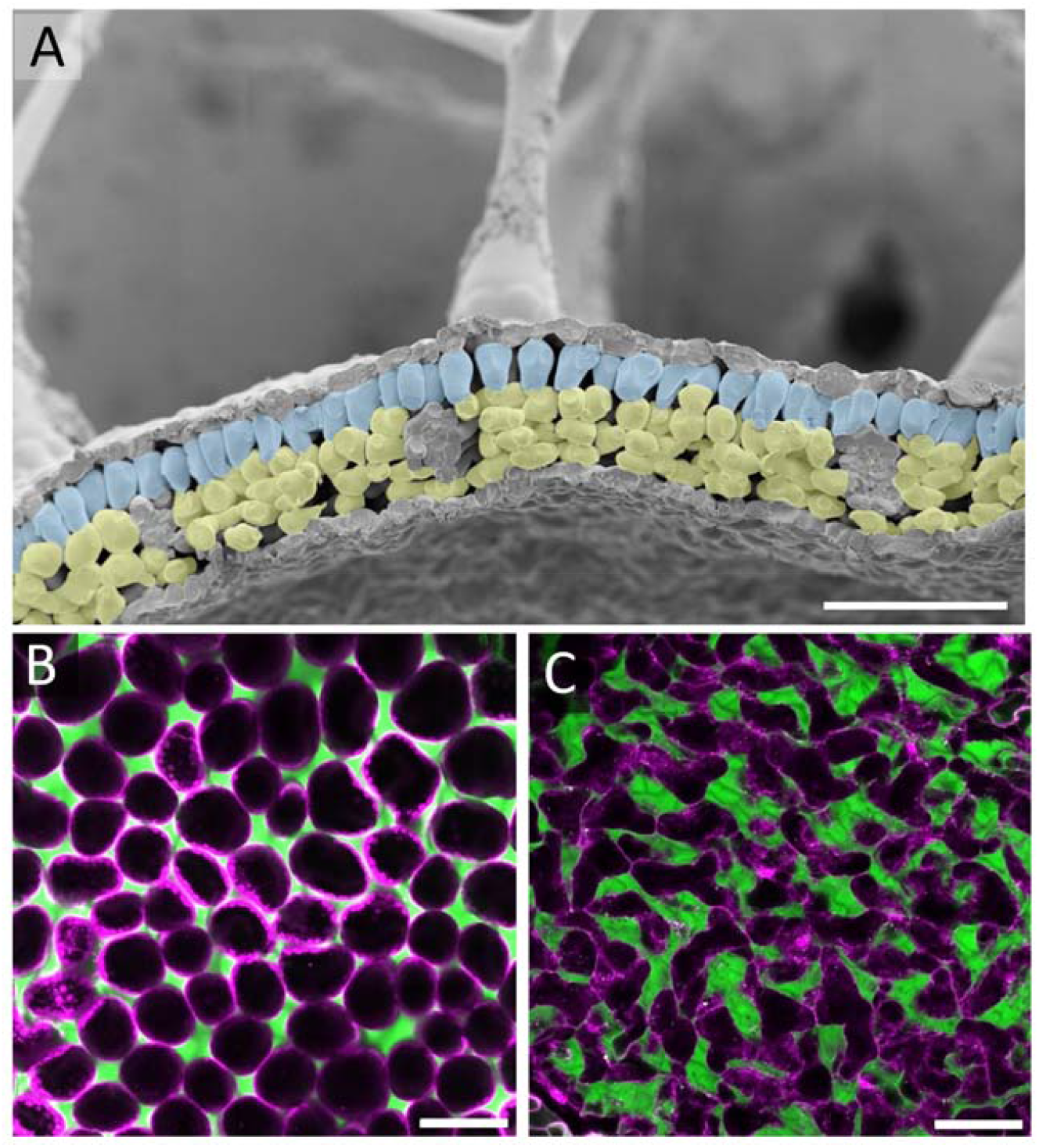
*A. thaliana* leaves contain palisade and spongy mesophyll cells surrounded by intercellular air space. (A) Transverse section of a mature *A. thaliana* leaf showing palisade mesophyll (false coloured blue) and spongy mesophyll (false coloured yellow) visualised by freeze-fracture cryoSEM. (B and C) Palisade (B) and spongy (C) mesophyll tissue visualised with Nile Red dye. Air spaces are green, cell outlines magenta. Scale bar = 100μm.

Air space formation and patterning has been proposed to be controlled either cell autonomously in mesophyll cells (by local loss of cell adhesion (Jeffree, Dale and Fry, 1986) and increased growth of cell walls adjacent to air spaces (expansigeny) (Zhang *et al*., 2021; Treado *et al*., 2022)), or non-cell autonomously by a mechanical mechanism where the epidermis grows faster than the underlying mesophyll and pulls mesophyll cells apart to form and expand air spaces (Avery, 1933). There is some support for both hypotheses: recent live imaging and computational modelling has shown that air spaces in the spongy mesophyll expand by expansigeny, supporting the cell autonomous hypothesis in spongy mesophyll cells (Zhang *et al*., 2021; Treado *et al*., 2022). Reduction of epidermal growth by expression of the cell-cycle inhibitor *KIP*-*RELATED PROTEIN 1* (*KRP1*) also decreases mesophyll porosity non-cell autonomously in the palisade, suggesting that epidermal growth may control air space patterning non cell-autonomously (Lehmeier *et al*., 2017). However, no molecular regulators of leaf air space patterning have yet been identified.

We recently showed that the plant hormone BR promotes epidermal growth in plant stems, and that removing this growth in a BR biosynthetic mutant of the aquatic plant *Utricularia gibba* (*Ugdwf4*) alters the internal structure of stems by mechanically constraining the expansion of internal tissue (Kelly-Bellow *et al*., 2023). Previous work has also shown that epidermal expression of the BR receptor *BRI1* is sufficient to rescue leaf size in the loss of function *bri1-116* mutant background (Savaldi-Goldstein, Peto and Chory, 2007), so it is possible that BR acts in a similar way in both leaves and stems. As the epidermis has been proposed to drive intercellular air space expansion (Avery, 1933), we hypothesised that BR promotes epidermal growth in leaves to drive air space formation and patterning.

Here we test this hypothesis by quantifying intercellular air spaces in the first leaf of *Arabidopsis thaliana* throughout development and analysing the effect of mutations in BR synthesis and perception. We show that loss of BR function does not affect air space formation, but causes air spaces to be gradually reduced in the palisade throughout leaf development. We further show that epidermal BR perception controls air space patterning non-cell autonomously, supporting a role for the epidermis in leaf air space development. Finally, we reduce epidermal growth by ectopic expression of the growth repressor *BIG BROTHER* (Disch *et al*., 2006), and show that this also reduces air space size. Overall, we propose that BR controls leaf air space patterning non-cell autonomously by promoting epidermal growth.

## Results

### Leaf air spaces in the palisade and spongy mesophyll undergo different developmental trajectories in *Arabidopsis*

To determine the developmental timeline for air space formation in leaves, we used Nile red dye to quantify porosity (percentage air space) of *Arabidopsis thaliana* leaf one throughout development (Kawase *et al*., 2015, Kawase *et al*., 2016). Nile red emits two separate wavelengths of light when bound to a membrane or free in solution, allowing mesophyll cell membranes and intercellular air spaces to be visualised simultaneously (fig. 2A). Leaf air spaces are first detectable by Nile red in the spongy and palisade mesophyll at 6 days after sowing (DAS) (fig. 2A). As Nile red dye marks air spaces by entering through stomata and stomata are not visibly open before 6DAS we were unable to assess the porosity of leaves before 6DAS using this method. To overcome this difficulty we used the *pUBQ1::2x-tdTomato-29-1* line (Shapiro *et al*., 2015) to visualise mesophyll cell outlines at 4 to 6 days (fig S1). This showed that limited air spaces are present at 5 DAS, with no air spaces visible at 4 DAS in either spongy or palisade mesophyll.

**Figure 2.**
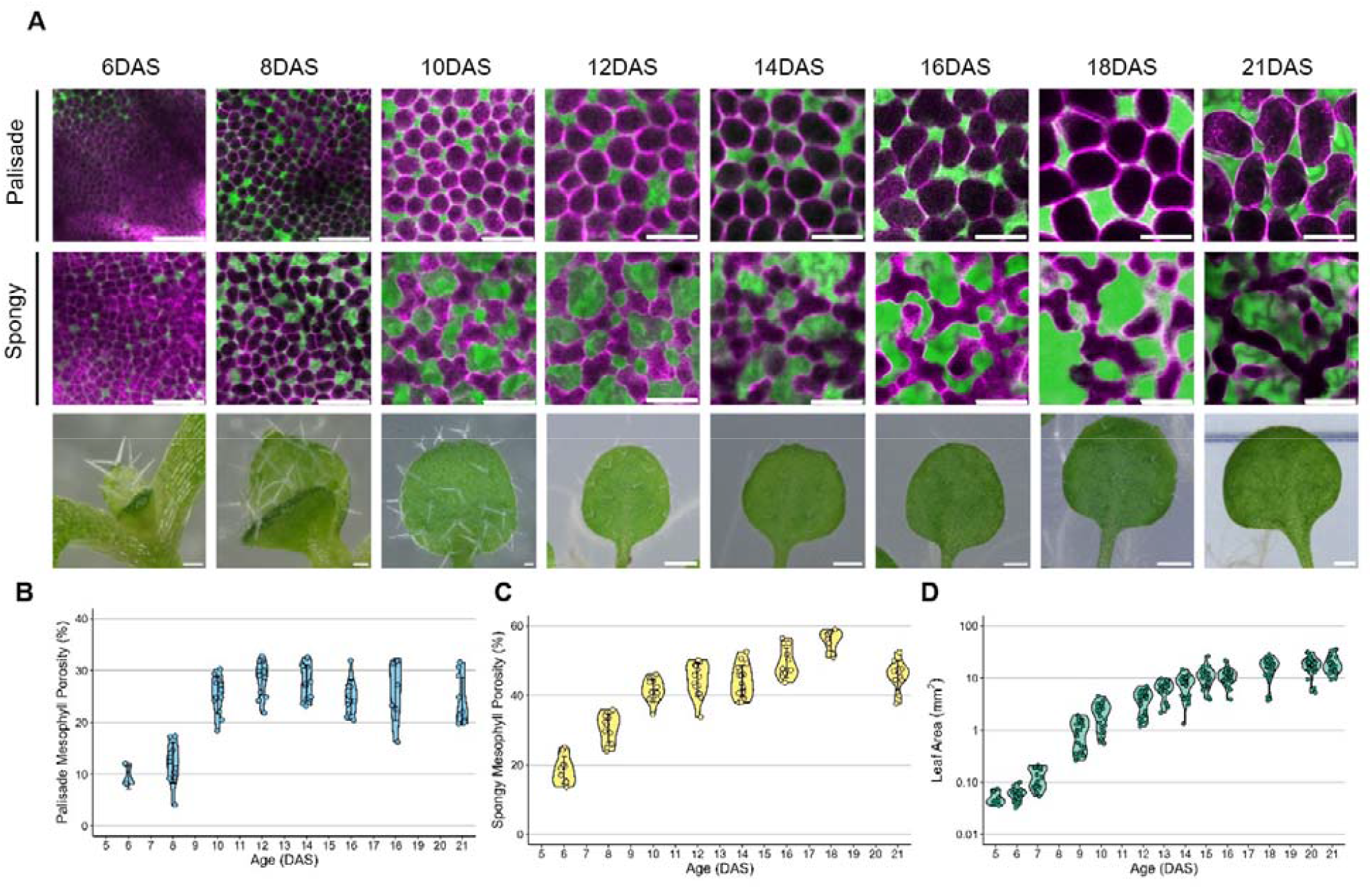
Palisade and spongy mesophyll air spaces undergo different developmental trajectories in *Arabidopsis thaliana*. (A) Air spaces in the Palisade (top row) and spongy (middle row) visualised by Nile red dye. Air spaces are green, cell outlines magenta. Bottom row of A shows image of whole leaf for scale. (B and C) Palisade (B) and spongy (C) mesophyll tissue porosity (%) of leaf 1 over 21 days. (D) Leaf area of *A. thaliana* leaf 1 over the same time period. Scale bar for (A): top two rows = 50μm; bottom row, 6-10 DAS = 100μm, 12-21 DAS = 1000μm

As leaf development proceeds in a basipetal gradient (Heynh *et al*., 1991; Donnelly *et al*., 1999; Fox *et al*., 2018) we quantified leaf porosity at the midpoint of the leaf blade to better compare across timepoints (fig. 2B and C). We observed markedly different trajectories for air space expansion in the palisade and spongy mesophyll. At 6 DAS, box shaped palisade mesophyll cells are mostly connected, with some small air spaces appearing at multicellular junctions (10% porosity, fig. 2A and B), whereas spongy mesophyll porosity is already 20% (fig. 2C). From 6 DAS to 21 DAS, spongy mesophyll porosity increased from 20% to 55%, with an initially faster rate that gradually slowed down until plateauing at around 18 DAS. Over this same period, palisade mesophyll porosity increased more rapidly from 10% at 8 DAS to around 30% at 10DAS, followed by no further increase despite leaf area continuing to expand (fig. 2B and D). Together, these results highlight the different trajectories for air space development in the spongy and palisade mesophyll, and show that air space expansion mainly occurs between 8DAS and 10DAS in the palisade.

### Brassinosteroid signalling is required for air spaces in mature leaves

We have previously shown that BR promotes growth in the epidermis of stems (Kelly-Bellow *et al*., 2023), and epidermal growth has been proposed to drive air space expansion (Avery, 1933). Therefore, we hypothesised that BR may drive leaf air space expansion by promoting epidermal growth. To test if BR regulates air space patterning, we quantified tissue porosity in mature leaves of plants with reduced BR synthesis (*dwf4* mutants or plants treated with the BR synthesis inhibitor bortezomib (BZR)) or perception (the BR receptor mutant *bri1-116*).

In mature 3-week old first leaves of *dwf4, bri1-116* and BZR-treated plants, air spaces were significantly reduced in the palisade mesophyll from around 25% to around 10% (Fig. 3A and B; P < 0.0002). Tissue porosity was also slightly but significantly reduced in the spongy mesophyll in *bri1-116* (from 45 to 35% porosity, P = 0.0000153), whereas porosity in *dwf4* mutants and plants treated with 5μM BZR were not significantly different from wild type (Fig. 3A and C). These data suggest that BR is necessary for either air space initiation or expansion in the palisade mesophyll, with a minimal role in the spongy mesophyll.

**Figure 3.**
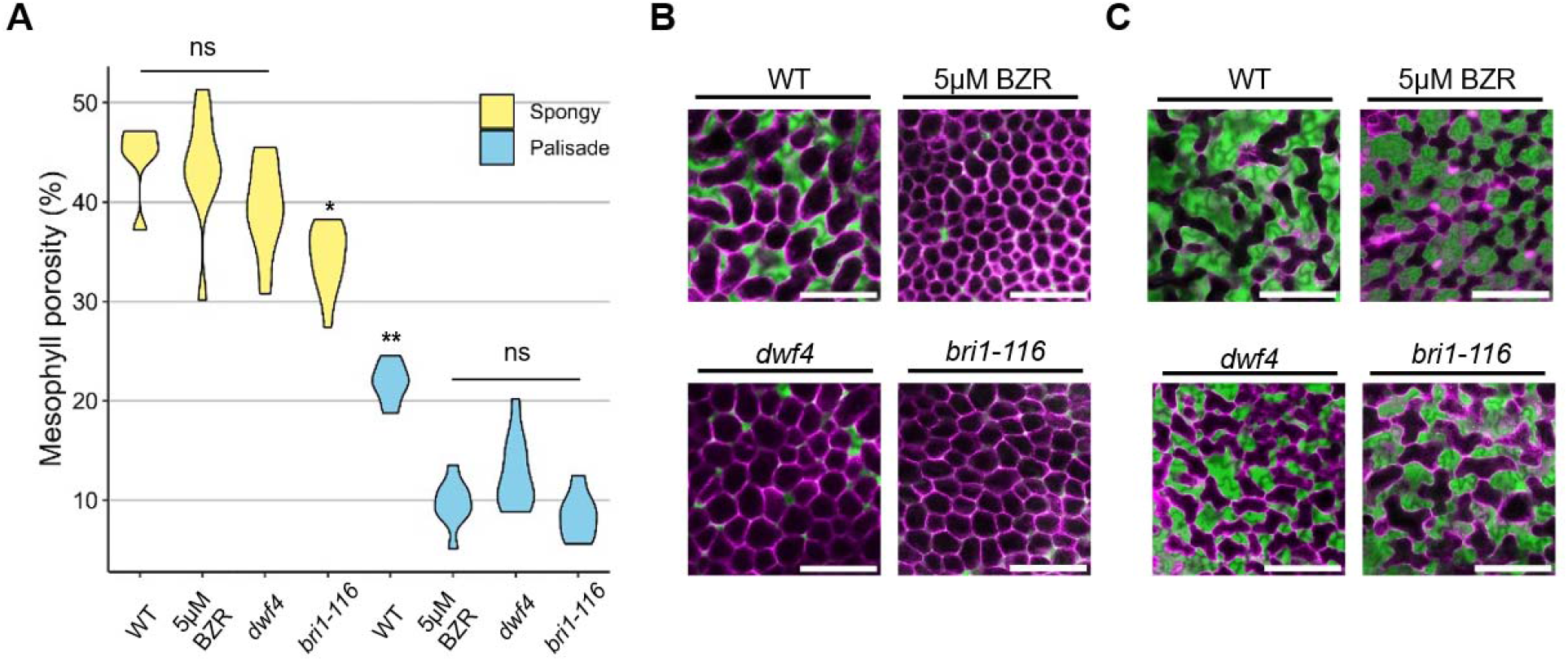
Brassinosteroids are required for normal air space patterning in leaves. (A) Porosity quantified using Nile red dye from sub-epidermal palisade and spongy mesophyll tissue layers at 21DAS. Plants were untreated wild-type (WT), wild-type plants grown in media containing 5μM Brassinazole (BZR), *dwf4* and *bri1-116* mutants. **(B)** Representative Nile red stained palisade and **(C)** spongy mesophyll images, with air spaces visible in green and cell outlines in magenta. (Scale bar, 100μm.) Two-way ANOVA; * = P < 0.05, ** = P > 0.01. ns= not significantly different.

### The BR perception mutant *bri1-116* forms air spaces in early leaf development which are lost by maturity

To determine if air space formation or expansion is prevented by a lack of BR signalling we analysed air space patterning throughout leaf development in the *bri1-116* mutant by staining with Nile red (fig. 4A). As previously, palisade mesophyll air spaces were difficult to measure prior to 8DAS. In young leaves at 8DAS, palisade mesophyll porosity was not significantly different between wild-type and *bri1-116* (P = 0.9998917), suggesting that BR does not drive air space formation (Fig. 4B). From 8DAS to 12DAS, palisade porosity in both wild-type and *bri1-116* increased from 10% to approximately 20%. Following this, *bri1-116* palisade mesophyll air space gradually decreased until approximately 7% at 21DAS, whereas wild type palisade porosity increased to around 25%. Spongy mesophyll porosity increases similarly in both wild type and *bri1-116* to around 40% at 12DAS (P= 0.9990888) (fig. 4 A and C). After this point wild type spongy porosity continues to increase to around 45% at 21DAS whereas *bri1-116* plateaus, ending up slightly, but significantly, smaller than wild type at 21 DAS (P= 0.0362756, fig. 4C). Spongy mesophyll cell morphology also appears unaffected in bri1-116 (fig. 4A). These data show that lack of BR perception does not affect air space initiation, but strongly reduces air space expansion in the palisade, but not the spongy mesophyll.

**Figure 4.**
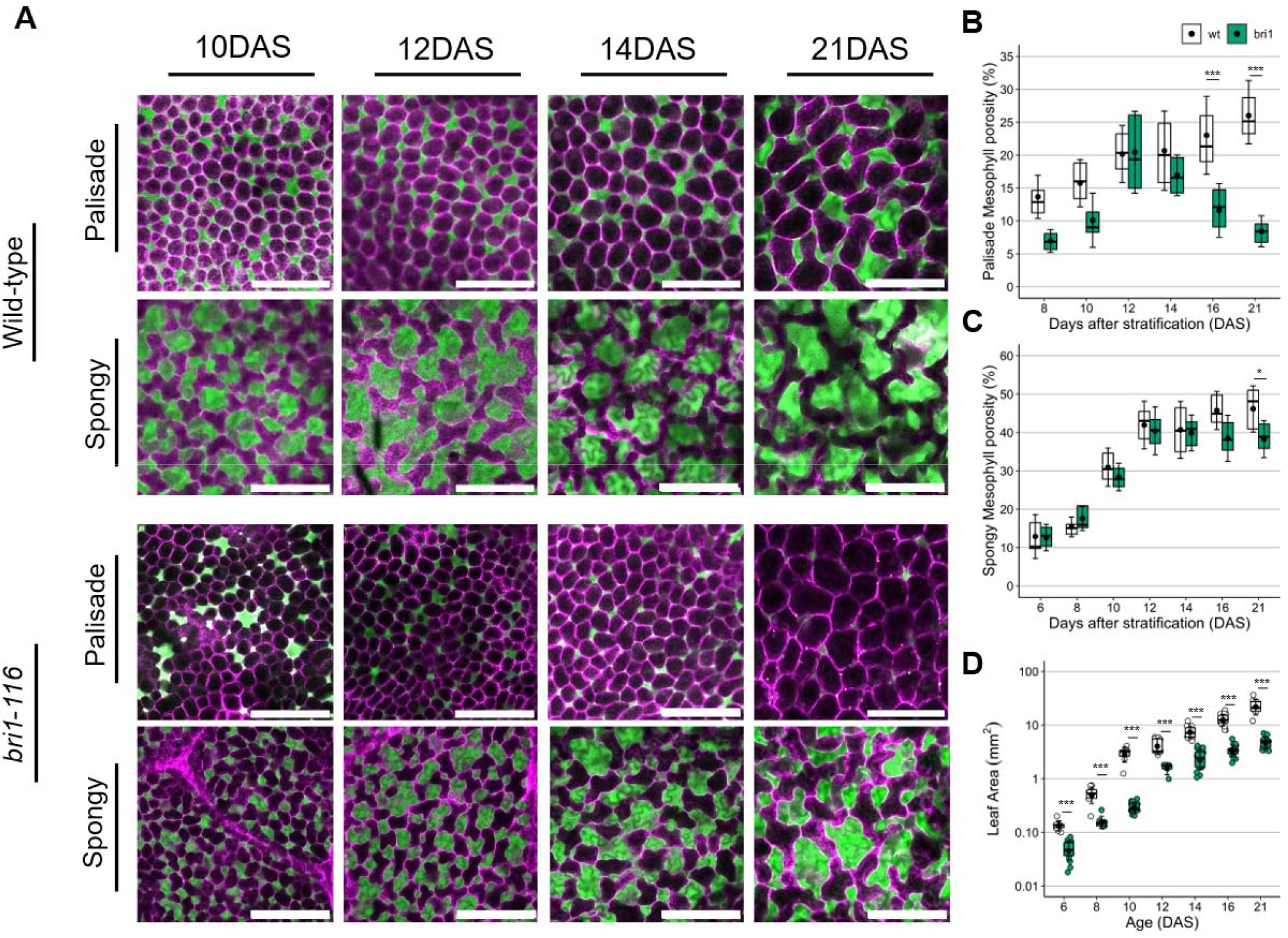
Loss of BR perception reduces palisade air space expansion but does not affect initiation. **(A)** Representative Nile red stained spongy and palisade mesophyll images from Col-0 and *bri1-116* throughout the first 21 days of leaf development. Air spaces are visible in green and cell outlines in magenta. (B and C) Palisade (B) and spongy (C) mesophyll porosity quantified over the same time period. (D) Leaf area quantified over 21 days of leaf development. (Scale bar, 20μm.) Two-way ANOVA; * = P < 0.05, *** = P > 0.001. Scale bar = 100μm.

### Epidermal brassinosteroid signalling is sufficient for air space expansion

One hypothesis to explain the initiation of palisade air spaces followed by their gradual reduction in size in the *bri1-116* mutant is that in wild type plants BR promotes epidermal growth, allowing internal tissue to grow with enough room for intercellular spaces to expand. Under this hypothesis, the *bri1-116* mutant exhibits reduced epidermal growth, but maintain normal mesophyll growth. This epidermal growth restriction constricts the expanding mesophyll tissue and presses palisade mesophyll cells together, resulting in a reduction of air space throughout development.

If the above hypothesis is correct, BR signalling in the epidermis should be sufficient to drive air space expansion in the palisade mesophyll. To determine if this is the case, we quantified the porosity of *bri1-116* mutants transformed with a *BRI1* tissue specific epidermal rescue line, *pATML1::BRI1-GFP* (Savaldi-Goldstein, Peto and Chory, 2007). As previously described, the epidermal rescue of *BRI1* signalling restored leaf shape and size and closely resembled a wild-type plant rosette (Fig. S2). Additionally, epidermal rescue of *BRI1* function rescued palisade mesophyll porosity to wild type levels (fig. 5A, P = 0.0795873). Overall, this suggests that BR signalling via the epidermis non-cell autonomously enables the retention of air spaces in the palisade mesophyll.

**Figure 5.**
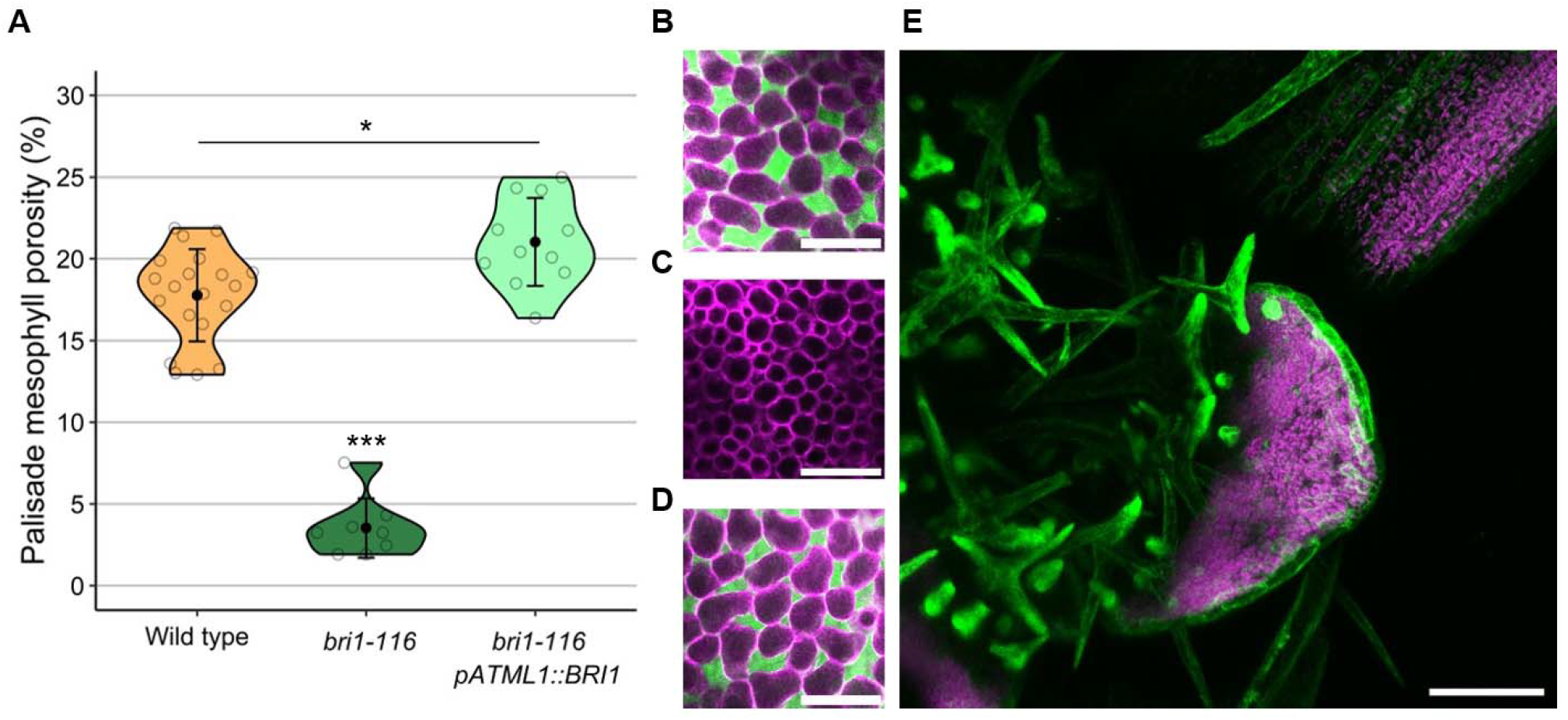
Epidermal expression of *BRI1* is sufficient to rescue mesophyll air space. (A) Quantification of palisade mesophyll porosity in leaf 1 & 2 of wild-type, *bri1-116* mutants and *bri1-116* mutants rescued with epidermal *BRI1-GFP* at 21DAS. (B-D) Palisade mesophyll of wild-type (B), *bri1-116* (C) and *bri1-116 pATML1::BRI1* rescue (D) leaves imaged with Nile red dye. (E) Leaf 2 of a *bri1-116 pATML1::BRI1-GFP* rescue at 7DAS. (B-E) Scale bar = 100μm. Kruskal-Wallis rank sums test; * = P < 0.05, *** = P > 0.001.

### Mesophyll cell division rate is unaltered in *bri1-116*

An alternative hypothesis to explain the reduced air space and increased cell density of the *bri1-116* mutant is that cell division in the mesophyll is increased, and that more cells are squashed into the same area. To test if this was the case we calculated palisade cell number per leaf in both wild type and *bri1-116* (fig. 6D). This showed no significant difference between wild type and *bri1-116*, despite *bri1-116* leaves being significantly smaller (P > 0.05). This suggests that division rates are broadly similar. To better quantify whether cell division is increased in *bri1-116* we performed clonal analysis by inducing Heatshock (HS)-inducible CRE-mediated lox site recombination via heatshock to induce GFP expression (fig. 6A) (Kuchen *et al*., 2012; Lee *et al*., 2019). We induced GFP expression in single cells and counted the number of cells in each clone two or three days later as a proxy for cell division rate. These data show that cell number per clone is not significantly different between wild type and *bri1-116* at two different timepoints across palisade development (8–10 and 14–17 days, P > 0.05) (Fig 6B, C, E and F). Overall, these data suggest that the decrease in palisade tissue porosity and increase in cell density observed in *bri1-116* is likely due to a restriction in epidermal growth while division rate in the palisade is maintained.

**Figure 6.**
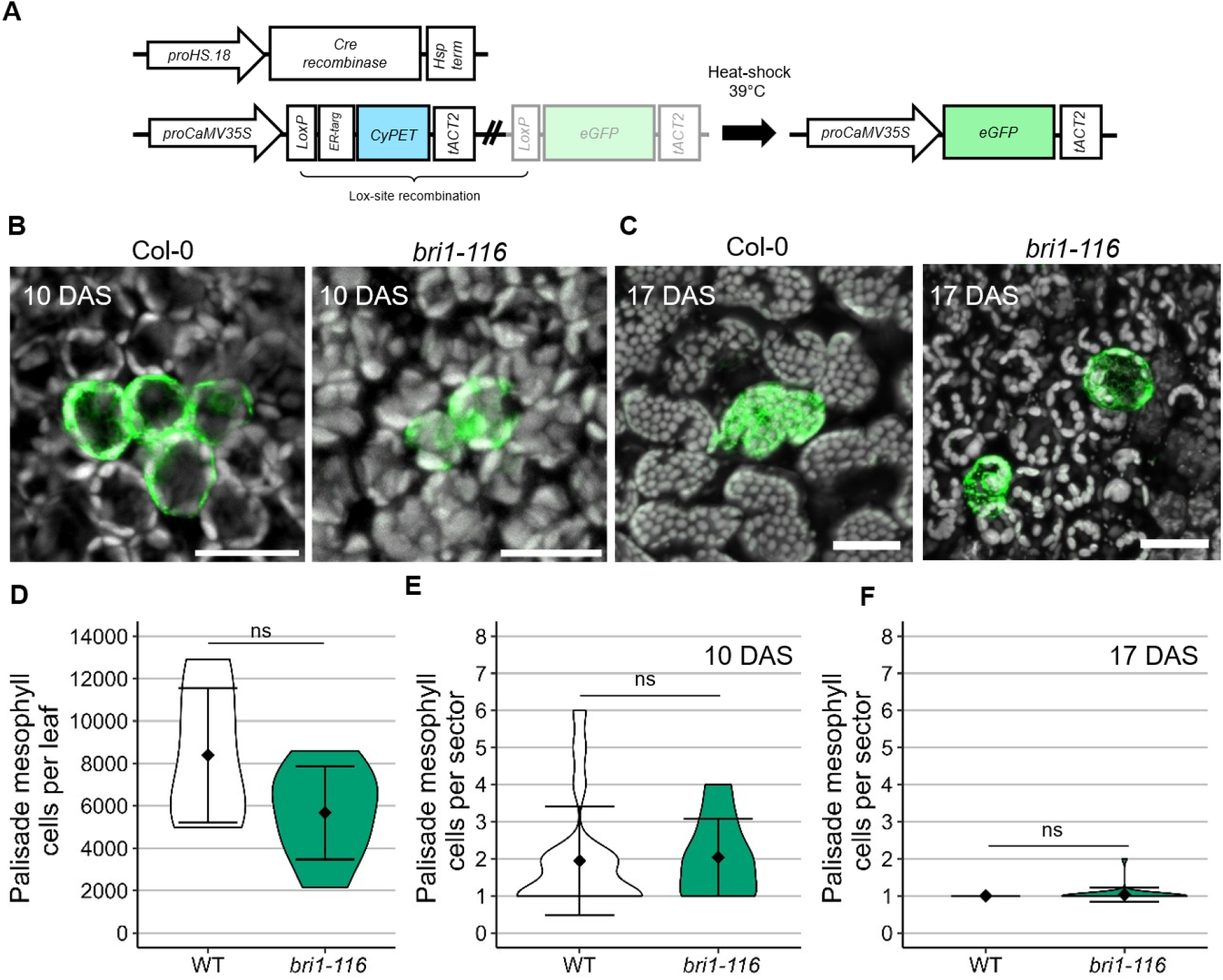
Palisade mesophyll cell division is not increased in BR signalling mutants. (A) Diagram of the heat-shock inducible CRE-lox recombinase system. (B and C) Wild-type and *bri1-116* palisade mesophyll sectors expressing GFP in (B) a 10 DAS leaf two days after sector induction and (C) in a 17 DAS leaf three days after sector induction. Chlorophyll autofluoresence visualised in grey, GFP in green. (D) Quantification of the number of palisade mesophyll cells in wild-type and *bri1-116* mutant leaves 1 & 2 at 21 DAS by extrapolation based on leaf area. (E and F) Quantification of cells in sectors expressing GFP in palisade mesophyll of 10 DAS leaves induced by heat-shock at 8 DAS (E) and 17 DAS leaves induced at 14 DAS. Scale bar = 20μm. Student’s t-test; ; * = P < 0.05.

### Decreased epidermal growth via tissue specific expression of *BIG BROTHER* decreases mesophyll air space

If the reduced porosity of palisade tissue of BR mutants is due to reduced epidermal growth constraining the expansion of inner tissue it should be possible to reduce palisade mesophyll porosity by reducing growth in the epidermis independently of BR. To do this we used HS-CRE-inducible CRE-lox recombination to induce expression of the negative growth regulator *BIG BROTHER (BB)* (Disch *et al*., 2006) under control of the *AtML1* promoter (fig. 7A) to reduce growth specifically in the epidermis (fig. S3). We induced epidermal BB expression at 3 DAS (before air spaces form) and quantified leaf size and porosity at 21 DAS when leaves were fully expanded. Leaves where epidermal *BB* expression was induced were reduced in area compared to uninduced and wild type controls (fig. 7B and C) and had significantly lower palisade porosity than wild type (10% in induced *AtML1::BB* vs 25% in WT (Fig. 7D and F, P ≤ 0.04). Spongy mesophyll porosity was also reduced in induced *AtML1::BB* leaves, but more moderately; 50% in induced AtML1::BB vs 60% in WT (fig. 7E and F). These data suggest that reducing epidermal growth can reduce air space expansion in the mesophyll tissue non-cell autonomously.

**Figure 7.**
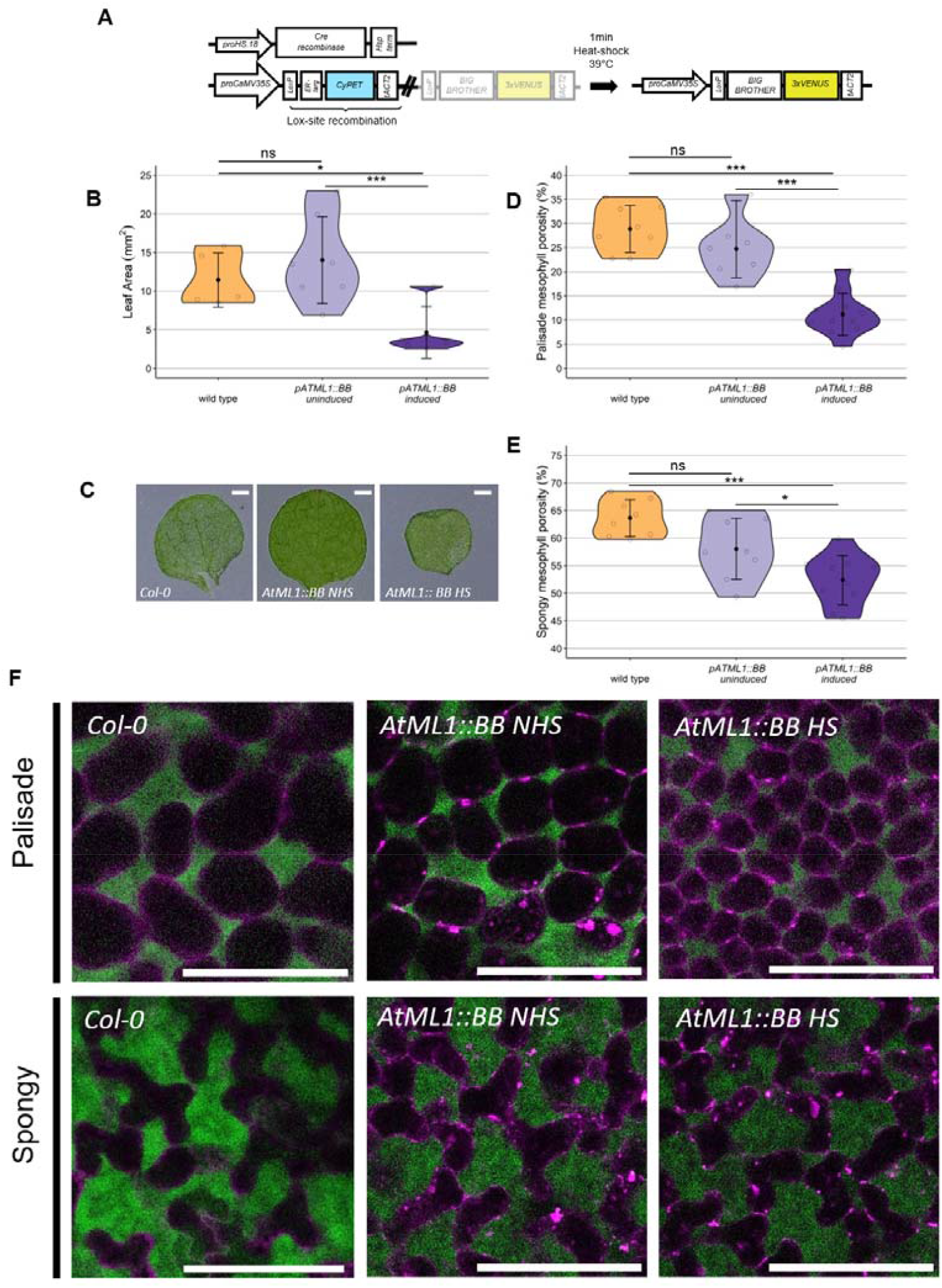
Repression of epidermal growth reduces palisade, but not spongy mesophyll porosity. (A) Construct design of heatshock-inducible epidermal *BIG BROTHER* expression. (B and C) Induction of epidermal *BB* at 3 DAS reduces leaf 1 area at 21 DAS. (D, E and F) Epidermal *BB* induction reduces palisade mesophyll porosity (D and F) but not spongy mesophyll porosity (E and F). Scale bars = 100μm in F and 1 mm in C. NHS = no heat shock, HS = heat shock. One-way ANOVA; ; * = P < 0.05, *** = P > 0.001.

## Discussion

Here we show that brassinosteroid signalling in the epidermis is necessary to pattern intercellular air spaces in leaf mesophyll tissue. Specifically, we show that BR is not required for air space initiation, but instead is required for retention and expansion of air spaces in the palisade mesophyll as the leaf expands. By experimentally reducing growth rate in the epidermis by epidermal-specific expression of the negative growth regulator *BIG BROTHER (BB)*, we also demonstrate that restricting epidermal growth is sufficient to reduce palisade mesophyll porosity. This is consistent with previous work showing that limiting cell division in the leaf epidermis leads to reduced palisade mesophyll porosity, but not spongy mesophyll porosity (Lehmeier *et al*., 2017). Therefore, we propose that brassinosteroids drive epidermal growth to determine the pattern of intercellular air spaces in the palisade mesophyll non-cell autonomously.

### Epidermal growth determines leaf air space pattering

Our data suggest that epidermal growth is necessary to allow intercellular air spaces to expand in the palisade mesophyll throughout leaf development. Previously, the epidermis has been proposed to drive air space formation and expansion by growing faster than the mesophyll cells and physically pulling them apart (Avery, 1933). Our data suggest that this is unlikely to be the mechanism driving initial air space formation (as lines with reduced epidermal growth still initiate air spaces), but it is possible that a faster growing epidermis acts to enhance air space expansion later in leaf development by applying tension to slower growing inner palisade tissue. Alternatively, epidermal growth may not drive air space expansion, but might simply need to be high enough not to mechanically constrain the palisade mesophyll tissue as it grows. It may be possible to distinguish these two hypotheses in the future by further experiments manipulating growth rates specifically in the mesophyll and epidermis.

These data contrast with the prevailing model of stem elongation, where faster growing internal tissue drives growth by putting a slower growing, stiffer, epidermis under tension. Instead, in the leaf the epidermis must grow sufficiently to allow (or drive, depending on the hypothesis) appropriate patterning of internal tissues. The fact that the non-cell autonomous effect of BR is mediated by the mechanical effect of differential growth between cell layers is also in line with recent work (Blanco-Touriñán *et al*., 2024) showing that BRI1 largely acts cell autonomously at the molecular level.

### Different developmental mechanisms control air space patterning in the spongy and palisade

Previous studies have described early spongy mesophyll development in detail (Zhang *et al*., 2021; Zhang and Ambrose, 2022), illustrating how air spaces first appear at multi-cell junctions after the degradation of the middle lamella, followed by intracellular turgor and microtubule patterning causing spongy mesophyll cells to change their shape from box-like to rounded to lobed. Consistent with previous work, our results indicate that both in the spongy and the palisade, air space initiation occurs at multicellular junctions (Sifton, 1945; Fox *et al*., 2018). However, our work shows that air space formation and expansion follow different trajectories in each mesophyll cell type: air spaces expand gradually throughout leaf development in the spongy, whereas the majority of air space expansion in palisade air space occurs between 8 DAS and 10 DAS (fig. 2). The period of high air space expansion in the palisade is coincident with the highest rate of leaf expansion, consistent with epidermal expansion having the largest effect on air spaces in this tissue layer.

BR also has a stronger effect on palisade air space development compared to spongy: palisade porosity in *bri1-116* strongly decreased from 12DAS onwards, whilst spongy porosity was less affected (fig. 4). Mature *bri1-116* spongy mesophyll cells also form complex lobed shapes similar to wild type (fig. 3C and 4A), suggesting that spongy mesophyll cell morphogenesis is controlled cell autonomously independently of BR.

### Interaction of BR with known mechanisms of air space patterning

It is currently unclear how the non-cell autonomous role of BR interacts with known regulators of air space patterning: stomata, chloroplasts and the microtubule cytoskeleton (Whitewoods, 2021).

Previous work has shown that BR represses stomatal formation and the *bri1-116* mutant has increased stomatal density (Kim *et al*., 2012). As stomata are associated with sub stomatal air chambers, they are generally positively correlated with palisade porosity (Lundgren *et al*., 2019). Therefore, the decreased porosity of the *bri1-116* mutant alongside increased stomatal density suggests that BR is necessary for stomata to promote air space formation. We hypothesise that epidermal growth acts in parallel to a stomatal signal and is necessary to allow substomatal cavities to expand. However, it is also possible that stomata signal via BR to locally promote air space formation.

Several mutants with defects in chloroplast biogenesis or metabolism are also known to have more extensive leaf air spaces (Li *et al*., 1995; González-Bayón *et al*., 2006). How BR interacts with chloroplasts to control air space patterning is currently unknown. We hypothesise that they are parallel pathways that regulate air space patterning independently. Double mutants or inhibiting BR synthesis chemically in chloroplast mutants will be necessary to test this.

Evidence from several studies (Zhang and Ambrose, 2022; Zhang *et al*., 2021) suggests that local reorientation of microtubules towards air space-adjacent cell walls controls spongy mesophyll lobe formation and air space patterning. As discussed above, lobe formation in spongy mesophyll cells appears unaffected in BR mutants. This is consistent with computational models of spongy mesophyll morphogenesis, which show that when tissue growth is constrained by an external force, spongy mesophyll cell lobe formation is unaffected but tissue porosity reduces (Treado *et al*., 2022). Together these data point to a model where cell autonomous and non-cell autonomous mechanisms work in parallel to generate the overall pattern of air space in a leaf.

### Conclusions

Overall, we show that BR controls leaf air space patterning non-cell autonomously by promoting epidermal growth. It remains to be seen whether this mechanism is simply a necessary prerequisite for air space expansion, or whether plants actively modulate epidermal BR signalling to alter air space patterning in response to environmental cues.

## Materials and Methods

### Plant Material and growth conditions

Plants were grown on plates containing MS media (0.441% Murashige and Skoog including vitamins, 0.05% (w/v) MES, 0.8% Difco agar, pH to 5.7). Sterilised seeds were stratified in the dark at 4°C for 2 days before growing in a controlled environment chamber at 20°C with 16 h of illumination (150 μmol m−2 s−1).

BZR (Sigma) was dissolved in water and added to plates to a final concentration of 5μM.

### Nile red staining and confocal imaging

Nile red dye (Sigma) was dissolved in 1.0 cs Silicon oil (Aston Chemicals) by shaking overnight to a final concentration of 10 μM. Leaves were stained by soaking in Nile red solution for 10 minutes before mounting on a slide under a coverslip and imaging on a Leica SP8 confocal microscope. Nile red bound to cell membranes was excited with a 561 nm wavelength laser and visualised between 565 and 640 nm, and Nile red free in solution was excited with a 488 nm wavelength laser and visualised between 490 and 560 nm. Unbound Nile red fluorescence was quantified as described previously (Kawase *et al*., 2015, Kawase *et al*., 2016) to calculate tissue porosity.

### Leaf area and cell counts

Whole plants were imaged using a Keyence VHX8000 microscope, and leaf area quantified using ImageJ. Cell counts per leaf were calculated by manually counting palisade cell number in a 200 × 200 micron region of palisade tissue from a leaf stained with Nile red. Cell number per leaf was calculated by extrapolating the cell number per unit area to the whole leaf area for each genotype.

### Construct generation

The GFP sector line contained two constructs: a p35S-loxPrev-CyPET-t35S-loxPrev-GFP-tAct sector construct (EC71924) and a pHSP18.2::CRE-tAtHS construct (EC71692). These were both generated by golden gate cloning as previously described (Weber *et al*., 2011). See genbank files (EC71924 and EC71692) in the supplementary data for final construct sequences, and Table S1 for Golden Gate module details. Plants were transformed with both constructs by floral dip and selected on plates using Kanamycin and Hygromycin selection. Homozygous T3 lines were crossed to *bri1-116* heterozygotes and F2 offspring homozygous for *bri1-116* and containing the sector constructs were used for sector analysis.

The Epidermal inducible BIG BROTHER line (AtML::BB) contained a pL2B-BAR-HS18.2:CRE-U5-Cre-pAtML1::loxP-CyPET-t35S-loxP-BB-3xVenus-tAct (CW0091). CW0091 was also generated by Goldengate cloning. See genbank files (CW0091) in the supplementary data for final construct sequence and Table S1 for Golden Gate module details.

### Sector analysis and heatshock induction

Plants of the GFP sector line were grown on plates and heat-shocked by floating in a 39°C water bath for 1-2 minutes. Heat-shocked plants were returned to the growth room and left to grow for 2 or 3 days before imaging GFP clones in leaf 1 using a Leica SP8 confocal microscope. Cell numbers per clone were manually counted in multiple sectors across 5 leaves.

Plants of the AtML1:BB line were grown on plates and heat-shocked at three days old by floating in a 39°C water bath for 12 minutes to induce constitutive CRE recombination and ectopic *BB* expression in the entire epidermis.

## Acknowledgements

We thank Ray Wightman for help with Scanning Electron Microscopy and Jean Dillon and Edwyn Locke Bevia for helpful comments on the manuscript.

## Funding

This work was funded by a Career Development Fellowship awarded to CW (award G110004) and a Cambridge School of Biological Science (SBS) DTP studentship awarded to JF (PTAG-094)

## Author contributions

JF and CW performed conceptualization, developed methodology, undertook experimental investigation and data analysis and visualisation as well as manuscript writing and supervision. CW also provided project administration. AR and RDF performed experimental investigation and data analysis, and SF created the EC71924 sector line (generated resources).

---

**Supplementary figure S1.**
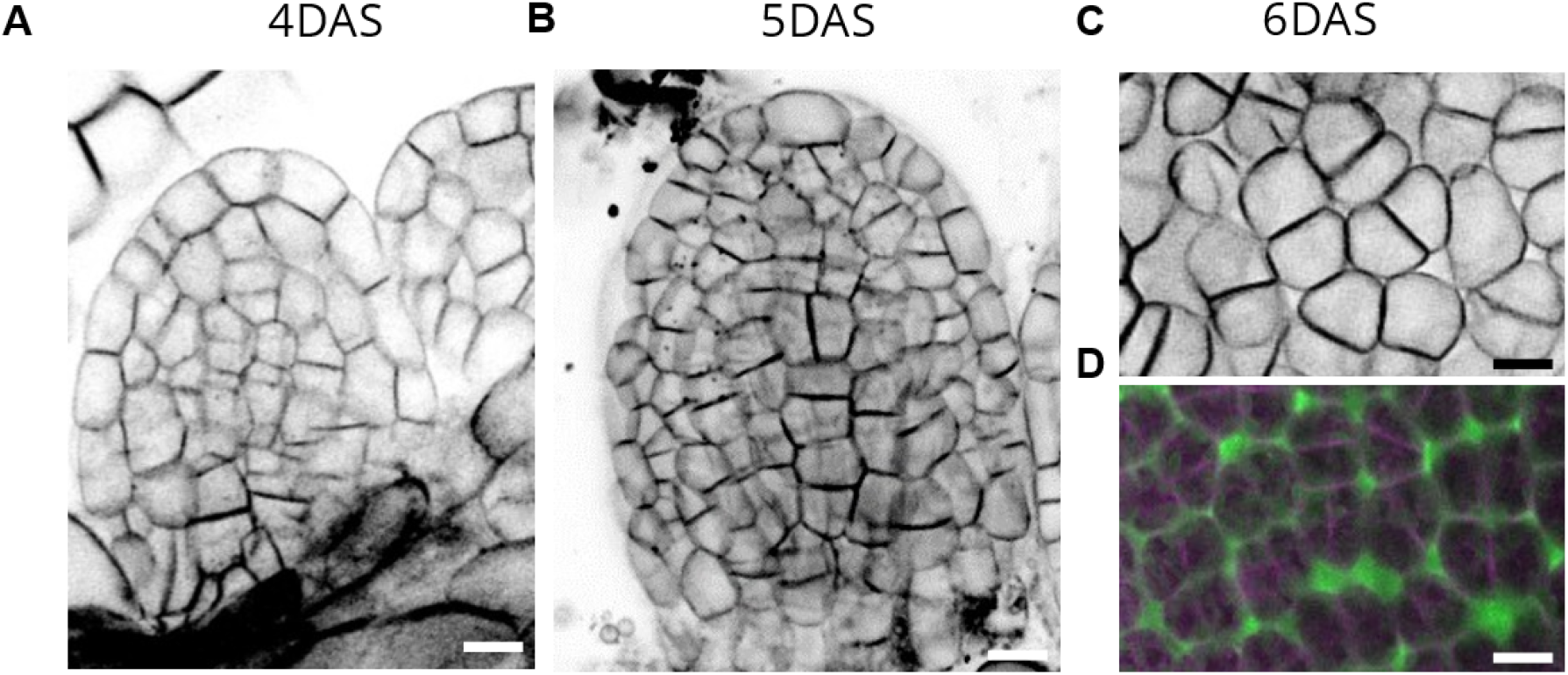
Air space formation happens around 5 DAS. (A-C) Confocal images of pUBQ1::2x-tdTomato-29-1 leaves from 4 – 6 DAS. (A) 4 DAS (B) 5 DAS and (C) 6 DAS. (D) Nile red dye staining of the abaxial side of leaf 2 at 6DAS. Scale bars = 10μm.

**Supplementary figure S2.**
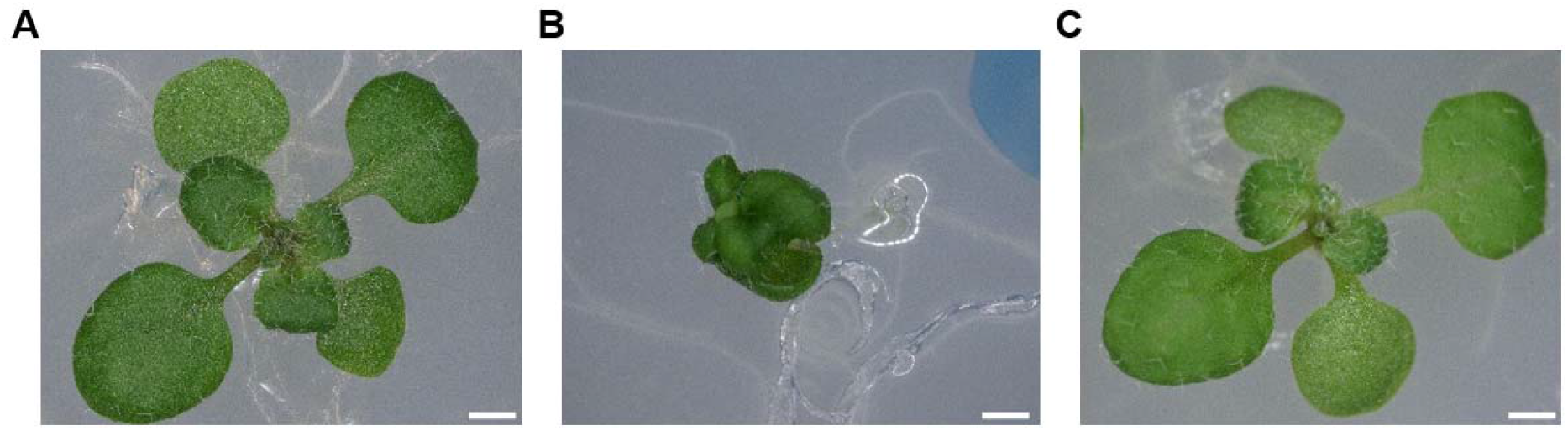
epidermal expression of *BRI1* rescues leaf size in *bri1-116*. (A-C) 14 day old *A. thaliana* seedlings; WT (A), *bri1-116* (B) or (C) *bri1-116 pATML1::BRI1-GFP*. Scale bars = 1mm.

**Supplementary figure S3.**
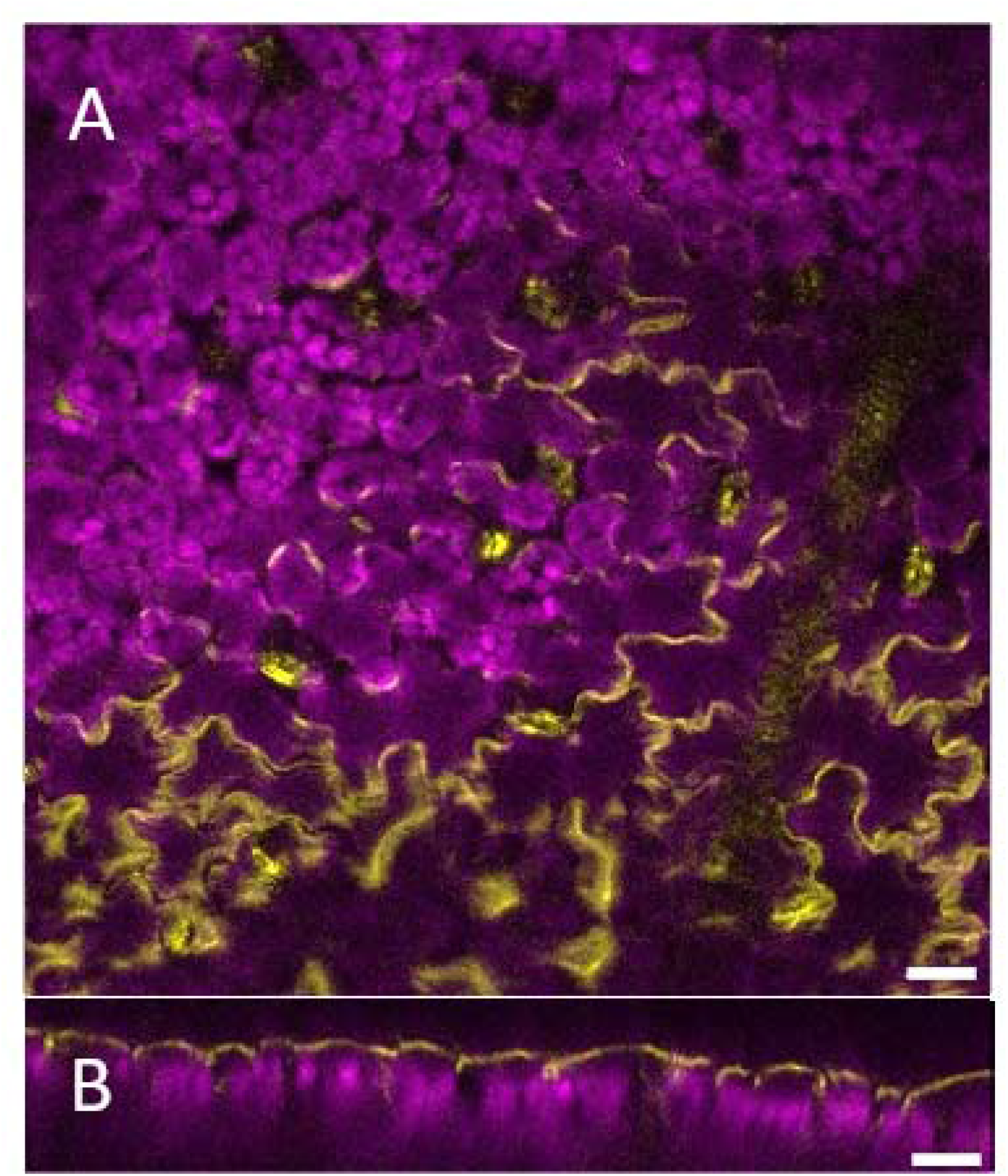
*AtML1::BB-3xVENUS* is induced specifically in the epidermis. (A) confocal image of a heatshock-induced 21 day old leaf where *BB-3xVENUS* expression was induced at 3 DAS. Palisade mesophyll cells lack VENUS expression, whereas epidermal cells show bright VENUS. VENUS fluorescence is yellow and Chlorophyll autofluorescence magenta. (B) Orthogonal view highlighting epidermal-specific expression of BB-3xVENUS. Scale bars = 100μm.

## Notes

### Competing Interest Statement

The authors have declared no competing interest.

